# Deficiency of the paternally inherited gene *Magel2* alters the development of separation-induced vocalization and maternal behavior in mice

**DOI:** 10.1101/2021.04.01.438102

**Authors:** Gabriela M. Bosque Ortiz, Gustavo M. Santana, Marcelo O. Dietrich

## Abstract

The behavior of offspring results from the combined expression of maternal and paternal genes. Genomic imprinting silences some genes in a parent-of-origin specific manner, a process that, among all animals, occurs only in mammals. How genomic imprinting affects the behavior of mammalian offspring, however, remains poorly understood. Here, we studied how the loss of the paternally inherited gene *Magel2* in mouse pups affects the emission of separation-induced ultrasonic vocalizations (USV). Using quantitative analysis of more than one hundred thousand USVs, we characterized the rate of vocalizations as well as their spectral features from postnatal days 6 to 12 (P6-P12), a critical phase of mouse development during which pups fully depend on the mother for survival. Our analyses show that *Magel2* deficient offspring emit separation-induced vocalizations at lower rates and with altered spectral features. We also show that dams display altered behavior towards their own *Magel2* deficient offspring. In a test to compare the retrieval of two pups, dams retrieve wildtype control pups first and faster than *Magel2* deficient offspring. These results suggest that the loss of *Magel2* impairs the expression of separation-induced vocalization in pups as well as maternal behavior, both of which support the pups’ growth and development.

## Introduction

For the normal development, mammalian offspring need copies from both maternal and paternal genomes. Some genes, however, are expressed in a parent-of-origin specific manner. In other words, some genes are always expressed when inherited from the mother and some genes are always expressed when inherited from the father^1-3^. The process that regulates the expression of genes in a parent-of-origin specific manner is called genomic imprinting^4^.

Genomic imprinting depends on epigenetic modifications of the genome. These modifications do not alter the sequence of the DNA but the chemical structure of the DNA, thereby leading to altered gene expression^1,4^. For example, an imprinted gene can be silenced in the maternal genome, and only the paternal allele will be expressed in the offspring (or vice-versa). Thus far, genomic imprinting has only been found in flowering plants and mammals^2,5-7^.

The adaptive value of genomic imprinting remains a matter of intense theorization. A prevailing theory on the effects of genomic imprinting, known as the kinship theory^5-7^, posits that maternal genes balance the energy investment of the mother between offspring survival and her own while paternal genes favor offspring survival alone. For example, the expression of paternal genes in the offspring would favor growth, while the expression of maternal genes would stunt growth^2,8-11^. The influence of imprinted genes is not restricted to growth, however. Imprinted genes are also primarily involved in brain development and in social behaviors^2,8-10^.

Consider, for example, a series of imprinted genes in human chromosome 15 that are paternally inherited^5–9^. The deletion of these paternally inherited genes in chromosome 15 leads to neurodevelopmental disorders, such as Prader-Willi syndrome (PWS). PWS presents with hypotonia and poor feeding early in life, followed by hyperphagia, alteration in social behavior, and cognitive deficits. It should be noted that because PWS involves several genes in human chromosome 15, it masks the relative contribution of single imprinted genes on the phenotype of the offspring. Among the PWS-related genes, MAGEL2 is a candidate gene for some of the clinical features of PWS. Humans with loss-of-function mutations in MAGEL2 present clinical aspects of PWS and autism spectrum disorders^10–12^, suggesting that this single paternally inherited gene supports at least some of the developmental alterations found in PWS. In agreement with the clinical features of MAGEL2 deficiency in humans, *Magel2* deficient mice show impairments in growth and adult social behaviors^10,13,14^. Despite these previous studies, a more systematic investigation of the effects of imprinted genes on behavior is necessary to understand the adaptive value of these genetic modifications in mammals.

We reasoned that investigating the effects of imprinted genes on offspring behavior during the early postnatal period—when the exchange of resources between the mother and the offspring is most important—can help shed light on the adaptive value of genomic imprinting. Towards this end, we studied how the loss of paternally inherited *Magel2* affects the vocal behavior of mouse pups when separated from their dams^15–23^, as separation-induced vocalizations signal the needs of the pups to the dams^19,24–29^. In contrast to human babies, mouse pups vocalize in the ultrasonic frequency range (45 - 120 kHz)^20,21,23,30^, which humans cannot hear. In order to survey the vocal behavior of mice, we recorded the emission of ultrasonic vocalizations (USVs) when pups were separated from the home nest. We performed these studies at postnatal days 6, 8, 10, and 12, since it is during this phase of mouse development that the peak expression of separation-induced vocalizations typically occurs^15,31,32^. We then used VocalMat^33^ to perform quantitative analysis of mouse vocal behavior. Moreover, we employed a maternal retrieval assay to test the effects of the loss of paternally inherited *Magel2* on maternal behavior towards the offspring.^12-17^ Our analysis shows that in mouse pups the deficiency of *Magel2* impairs the expression of separation-induced vocalizations. This deficiency also reduces maternal retrieval behavior towards *Magel2* deficient pups compared to non-deficient siblings.

## Results

### Early waning of vocal behavior in *Magel2*^m+/p−^ deficient pups

To investigate the effects of paternally inherited *Magel2* on the vocal behavior of infant mice, we crossed heterozygote males for *Magel2* deficiency with wildtype females. From this cross, we generated *Magel2* deficient offspring (*Magel2*^m+/p−^) that carry the null allele from the father (p−) and the imprinted allele from the mother (m+). This cross also generates wildtype littermates (*Magel2*^m+/p+^), used as experimental controls. As previously reported^15^, *Magel2*^m+/p−^ pups display lower body weight compared to controls (genotype: *F*_1, 181_ = 19.71, *P* < 10^−4^; age: *F*_3, 181_ = 69.64, *P* < 10^−10^; genotype x age: *F*_3, 181_ = 0.02, *P* = 0.99; two-way ANOVA; **Figure S1**). Post-hoc analysis shows a significant difference in body weight between genotypes at P12 (control: 6.08 ± 0.14 g; *Magel2*^m+/p−^: 5.51 ± 0.20 g; *P* = 0.04; Holm-Šídák test; **Figure S1**).

We recorded the emission of USVs during 20 minutes of separation from the home nest at different postnatal ages (P6, P8, P10, and P12; **Figure 1A**). First, we analyzed the total number of USVs emitted during the period of separation using two-way ANOVA. We found a significant effect of genotype, age, and interaction between genotype and age (genotype: *F*_1, 181_ = 20.61, *P* < 10^−4^; age: *F*_3, 181_ = 11.80, *P* < 10^−6^; genotype x age: *F*_3, 181_ = 3.90, *P* = 0.01; **Figure 1B**). Post-hoc analysis (Holm-Šídák test) shows that the total number of USVs is similar among groups at P6 (control: 991 ± 139 USVs; *Magel2*^m+/p−^: 787 ± 130 USVs; *P* = 0.75), P10 (control: 503 ± 45 USVs; *Magel2*^*m+/p−*^: 466 ± 70 USVs; *P* = 0.98), and P12 (control: 594 ± 71 USVs; *Magel2*^m+/p−^: 345 ± 46 USVs; *P* = 0.06) (**Figure 1B**). Compared to controls, however, *Magel2*^m+/p−^ pups show a ≈53% reduction in the emission of USVs at P8 (control: 1045 ± 100 USVs; *Magel2*^m+/p−^: 495 ± 65 USVs; *P* < 10^−4^). We also analyzed the data in 5-minute intervals and found similar effects of *Magel2* deficiency on the rate of vocalizations (**Figure S2**). Moreover, separating our analysis in females and males show similar effects of genotype and age (**Figure 1C-D** and **Table S1**). (Because we did not find differences in the rate of vocalization, we have pooled males and females in all further analysis). In sum, our results thus far show age-specific reductions in the emission of USVs in *Magel2*^m+/p−^ mice, suggesting a non-sex specific effect for paternally inherited *Magel2* on the vocal behavior of the offspring.

**Figure 1.**
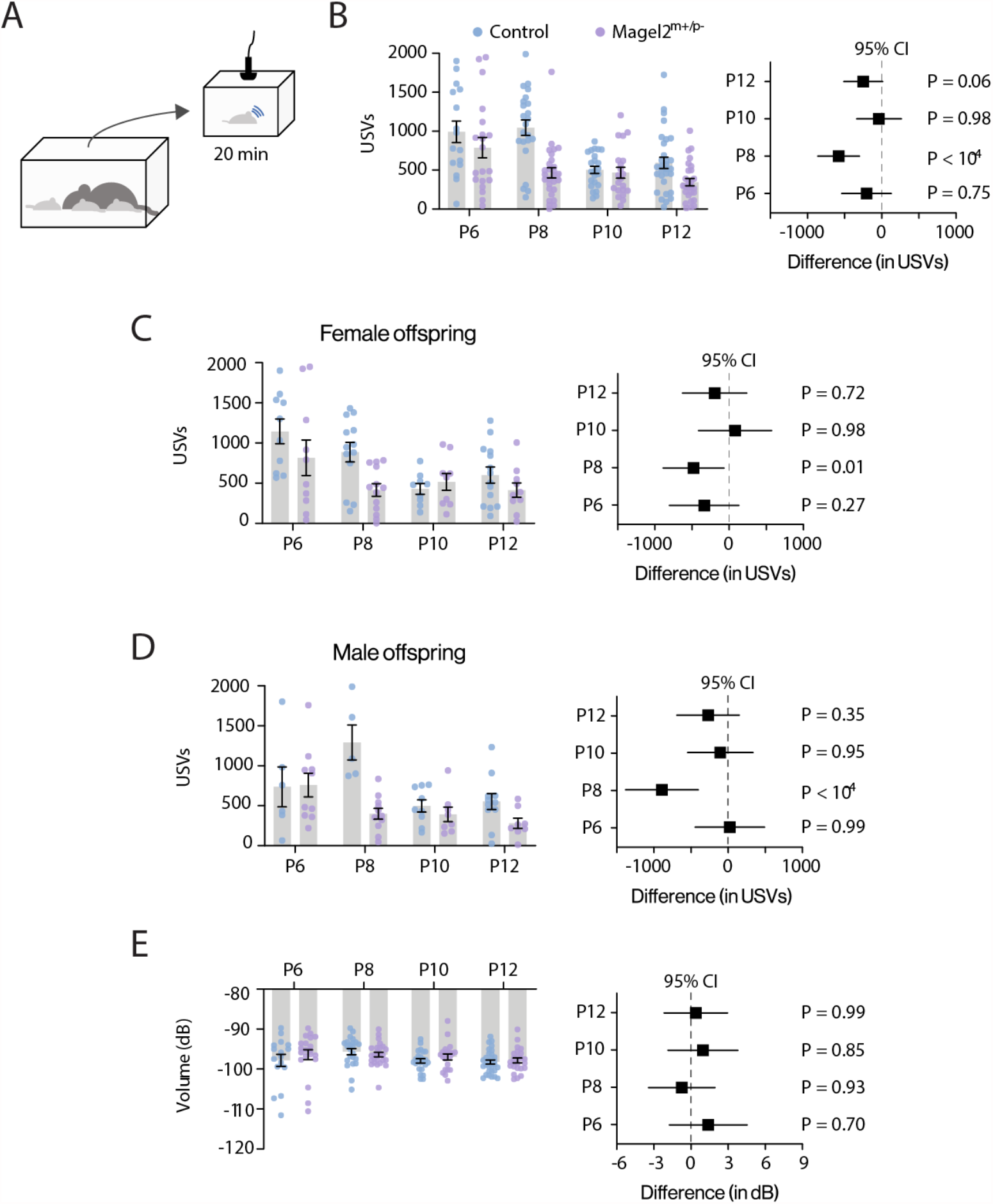
*Magel2* deficiency affects the emission of ultrasonic vocalizations in mice. (**A**) Schematic of the protocol used to record separation-induced USVs in mice (from P6 to P12); pups are separated from the home nest in a new chamber equipped with an ultrasonic microphone and recorded for 20 minutes. (**B**) Total number of USVs emitted by control (blue) and *Magel2* deficient (purple) littermates at P6, P8, P10, and P12; right panel denotes the 95% confidence intervals as a measure of effect size. (**C**) Similar to (B), but only considering female pups. (**D**) Similar to (B), but only considering male pups. (**E**) Average intensity of the USVs measured in decibels; right panel denotes the 95% confidence intervals as a measure of effect size. Bars represent mean value with error bars representing SEM and round symbols representing individual values. When plotting the effect sizes, squared symbols and black lines represent 95% confidence intervals calculated as the different between *Magel2* deficient and control pups. *P* values are provided in the figures as calculated using Sidak’s multiple comparison test. The sample sizes for control and *Magel2* deficient pups are: P6, n = 16 and 20; P8, n = 23 and 28; P10, n = 23 and 20; and P12, n = 30 and 28, respectively.

The emission of USVs occurs when the breathing musculature contracts, expelling air from the lungs and propelling it through the larynx^12,18^. Since previous reports found that *Magel2*^m+/p−^ mice display hypotonia^19^, we considered the hypothesis that the low rate of USV emission in *Magel2* deficient pups is due to a lower capacity to expel air from the lungs. To rule out this hypothesis, we measured the intensity (or volume, in decibels) of the USVs. Since the intensity of the USVs relates to the pressure by which the air is expelled through the larynx^18^, a lower intensity is expected in cases of hypotonia. This analysis shows that the intensity of the emitted USVs between *Magel2*^m+/p−^ mice and controls is similar in all ages tested (genotype: *F*_1, 181_ = 2.67, *P* = 0.10; age: *F*_3, 181_ = 2.33, *P* = 0.08; genotype x age: *F*_3, 181_ = 0.78, *P* = 0.51; two-way ANOVA; **Figure 1E**). Thus, hypotonia does not seem to be a factor of primary significance for the lower rate of USV emission in *Magel2*^m+/p−^ mice^16^.

### *Magel2*^m+/p−^ mice emit vocalizations with distinct spectral features

In addition to the rate of separation-induced vocalizations, the spectral features of the USVs also correlate with altered maternal care^20^. To test the extent to which *Magel2* deficiency affects the spectral features of USVs across ages, we used two-way ANOVA to analyze the frequency characteristics (pitch) and duration (**Figure 2A**) of USVs^21^. We found significant effects of genotype and age for maximal frequency (genotype: *F*_1, 181_ = 20.51, *P* < 10^−4^; age: *F*_3, 181_ = 3.23, *P* = 0.02; genotype x age: *F*_3, 181_ = 6.18, *P* < 10^−3^; **Figure 2B**), bandwidth (genotype: *F*_1, 181_ = 9.91, *P* < 10^−2^; age: *F*_3, 181_ = 6.97, *P* < 10^−3^; genotype x age: *F*_3, 181_ = 5.92, *P* < 10^−3^; **Figure 2E**), and duration (genotype: *F*_1, 181_ = 4.08, *P* = 0.04; age: *F*_3, 181_ = 6.35, *P* < 10^−3^; genotype x age: *F*_3,181_ = 2.05, *P* = 0.11; **Figure 2F**). In addition, we found a significant effect of genotype for mean frequency (genotype: *F*_1, 181_ = 4.86, *P* = 0.02; age: *F*_3, 181_ = 0.24, *P* = 0.87; genotype x age: *F*_3, 181_= 0.41, *P* = 0.74; **Figure 2D**), but not for minimal frequency (genotype: *F*_1, 181_ = 0.47, *P* = 0.47; age: *F*_3, 181_ = 1.96, *P* = 0.12; genotype x age: *F*_3, 181_ = 0.48, *P* = 0.69; **Figure 2C**). We then used post-hoc analysis (Holm-Šídák test) and found that at P8—but not at P6, P10, or P12— *Magel2*^m+/p−^ mice show significant differences compared to control mice in maximal frequency, bandwidth, and duration (see panels plotting the 95% confidence intervals in **Figure 2**).

**Figure 2.**
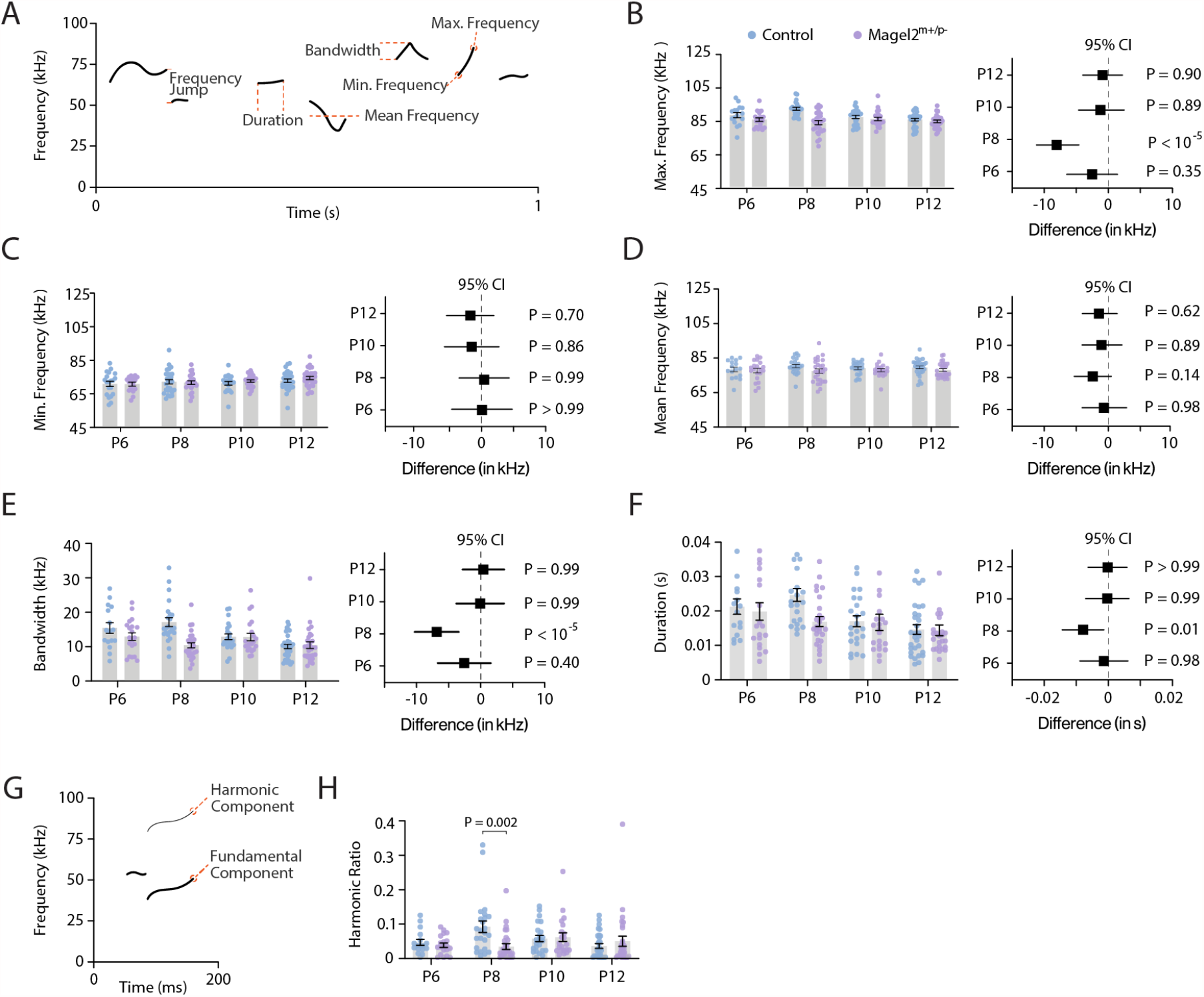
*Magel2* deficient pups emit ultrasonic vocalizations of distinct spectro-temporal features. (**A**) Illustration of a spectrogram with the spectro-temporal features measured for each USV. (**B**) Maximum frequency of the USVs emitted by control and *Magel2* deficient littermates at P6, P8, P10, and P12; right panel denotes the 95% confidence intervals as a measure of effect size. (**C**) Similar to (B) but plotting the minimum frequency of the USVs. (**D**) Similar to (B) but plotting the mean frequency of the USVs. (**E**) Similar to (B) but plotting the bandwidth of the USVs. (**F**) Similar to (B) but plotting the duration of the USVs. (**G**) Illustration of the spectrogram of a single USV with a harmonic component. (**H**) Ratio of harmonic across all USVs emitted by control and *Magel2* deficient littermates. Bars represent mean value with error bars representing SEM and round symbols representing individual values. When plotting the effect sizes, squared symbols and black lines represent 95% confidence intervals calculated as the different between *Magel2* deficient and control pups. In B-F, *P* values are provided in the figures as calculated using Sidak’s multiple comparison test post hoc analysis from two-way ANOVA test. In H, *P* values are provided as calculated using Mann-Whitney test. The sample sizes for control and *Magel2* deficient pups are: P6, n = 16 and 20; P8, n = 23 and 28; P10, n = 23 and 20; and P12, n = 30 and 28, respectively.

In addition to the main frequency component, USVs can contain harmonics (**Figure 2G**). We calculated the percentage of USVs with harmonics and found a significantly lower number in *Magel2*^m+/p−^ mice compared to controls at P8 (control: 9.0 ± 1.7 %; *Magel2*^m+/p−^: 3.5 ± 0.8 %; *U* = 145, *P*_2-tailed_ = 0.002, Mann-Whitney test; **Figure 2H**) but not at P6 (control: 4.7 ± 0.9 %; *Magel2*^m+/p−^: 3.8 ± 0.6 %; *U* = 142.5, *P*_2-tailed_ = 0.93, Mann-Whitney test; **Figure 2H**), P10 (control: 5.8 ± 0.9 %; *Magel2*^m+/p−^: 6.1 ± 1.2 %; *U* = 240, *P*_2-tailed_ = 0.98, Mann-Whitney test; **Figure 2H**), or P12 (control: 3.6 ± 0.7 %; *Magel2*^m+/p−^: 5.0 ± 1.5 %; *U* = 408.5, *P*_2-tailed_ = 0.98, Mann-Whitney test; **Figure 2H**). In sum, these results suggest that the loss of paternally inherited *Magel2* in mice causes discrete changes in the features of separation-induced vocalizations that are most evident at postnatal day eight.

### Discrete changes in the use of syllable types by *Magel2*^m+/p−^ mice

Mouse pups emit USVs of distinct classes—i.e., syllable types. Thus, the emission of different syllable types could explain the discrete changes in the spectro-temporal features of USVs in *Magel2*^m+/p− 21,22^. We used a validated software to automatically categorize each USV into one of eleven syllable types based on the morphology of the main component of the vocalization (**Figure 3A-B**) ^21^. The output of the method was the probability for each USV to be of a certain syllable type. The highest probability (*P*_1_) defined the syllable type for a given USV (**Figure 3A**).

**Figure 3.**
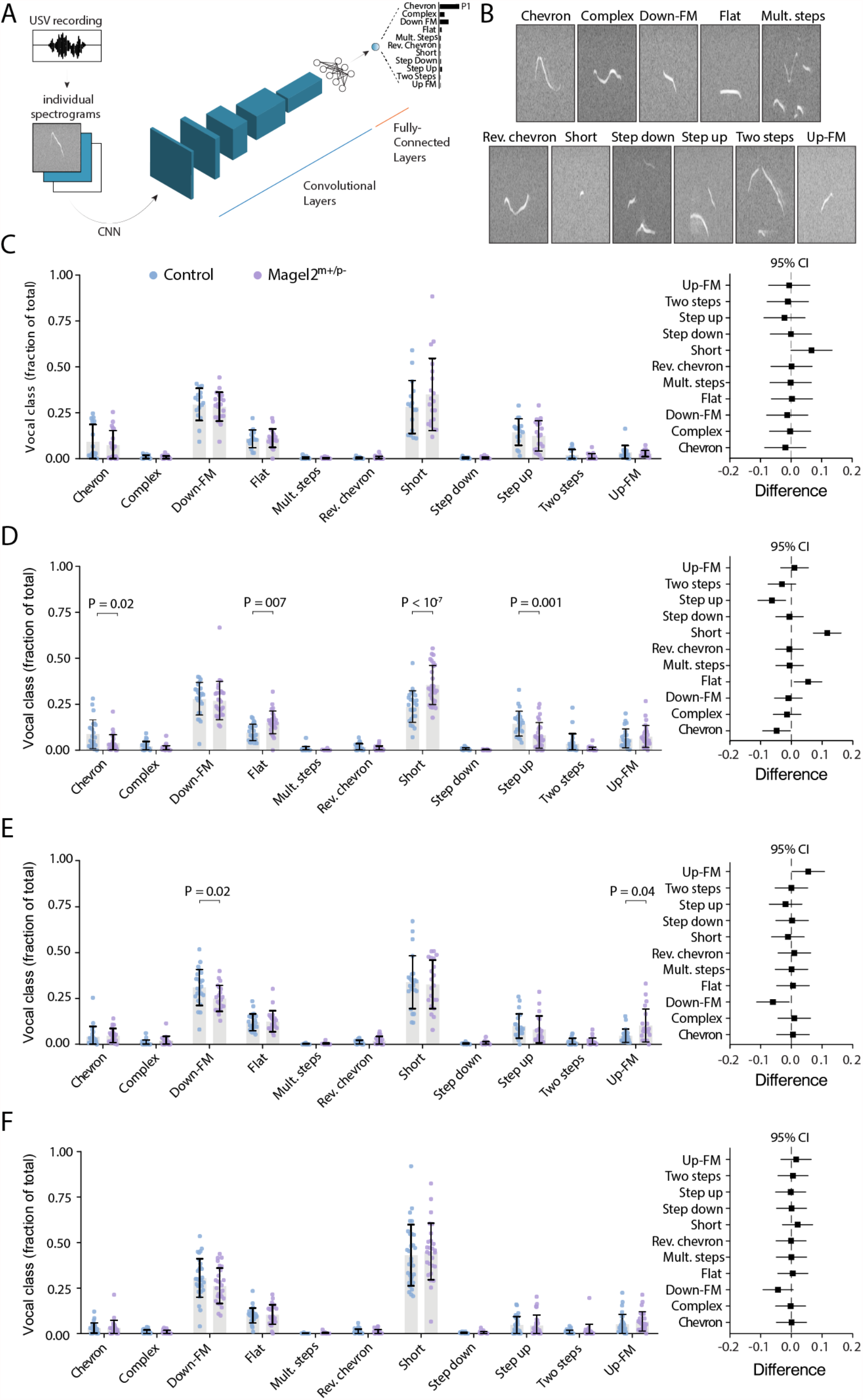
*Magel2* deficient pups emit ultrasonic vocalizations with discrete changes in the distribution of syllable types. (**A**) Illustration of the convolutional neural network used to classify each USV into one of eleven syllable types based on their morphology in spectrograms. (**B**) Spectrograms representing each of the eleven syllable types. (**C**) Distribution of syllable types in P6 pups—control in blue and *Magel2* deficient in purple. Data are showed as fraction of the total number of USVs; right panel denotes the 95% confidence intervals as a measure of effect size. (**D**) Similar to (C), but for P8 pups. (**E**) Similar to (C), but for P10 pups. (**F**) Similar to (C), but for P12 pups. Bars represent mean value with error bars representing SEM and round symbols representing individual values. When plotting the effect sizes, squared symbols and black lines represent 95% confidence intervals calculated as the different between *Magel2* deficient and control pups. *P* values are provided in the figures as calculated using Sidak’s multiple comparison test as a post hoc analysis after two-way ANOVA. The sample sizes for control and *Magel2* deficient pups are: P6, n = 16 and 20; P8, n = 23 and 28; P10, n = 23 and 20; and P12, n = 30 and 28, respectively.

The percent use of each syllable type per recording was compared between genotypes at each age using two-way ANOVA. Using this approach, we did not find any significant differences in the distribution of syllable types emitted by control and *Magel2*^m+/p−^ mice at P6 (genotype: *F*_1, 374_= 2.5 × 10^−16^, *P* > 0.99; class: *F*_10, 374_ = 93.25, *P* < 10^−4^; genotype x class: *F*_10, 374_ = 1.00, *P* = 0.44) and P12 (genotype: *F*_1, 616_ = 7.7 × 10^−16^, *P* > 0.99; class: *F*_10, 616_ = 278, *P* < 10^−4^; genotype x class: *F*_10, 616_ = 0.81, *P* = 0.62; **Figure 3C, 3F**). At P8, however, *Magel2*^m+/p−^ pups emit vocalizations of different syllable types (genotype: *F*_1, 539_ = 2.2 × 10^−15^, *P* > 0.99; class: *F*_10, 539_ = 167, *P* < 10^−6^; genotype x class: *F*_10, 539_ = 9.51, *P* < 10^−6^; **Figure 2D)**, which include: 55% less USVs of the type *chevron* (control: 9.0 ± 1.6 %; *Magel2*^m+/p−^: 4.0 ± 0.01 %; *P* = 0.02, Holm-Šídák test); 44% less USVs of the type *step-up* (control: 14.3 ± 1.4 %; *Magel2*^m+/p−^: 8.1 ± 1.3 %; *P* < 10^−3^, Holm-Šídák test); 36% more USVs of the type *flat* (control: 9.6 ± 0.9 %; *Magel2*^m+/p−^: 15.2 ± 1.2 %; P = 0.007, Holm-Šídák test); and 33% more USVs of the type *short* compared to controls (control: 23.6 ± 1.8 %; *Magel2*^m+/p−^: 35.5 ± 2.0 %; *P* < 10^−4^, Holm-Šídák test; **Figure 3D)**. At P10, we also identified discrete differences in the emission of USVs of different syllable types between groups (genotype: *F*_1, 462_ = 8.1 × 10^−17^, *P* > 0.99; class: *F*_10, 462_ = 149, *P* < 10^−4^; genotype x class: *F*_10, 462_ = 2.11, *P* = 0.02; **Figure 2E**): *Magel2*^m+/p−^ pups emit 19% less *down frequency modulation* (control: 31.0 ± 2.1 %; *Magel2*^m+/p−^: 25.0 ± 1.5 %; *P* = 0.02; Holm-Šídák test) and emit 53% more *up frequency modulation* (control: 4.8 ± 0.8 %; *Magel2*^m+/p−^: 10.3 ± 2.0 %; *P* = 0.04; Holm-Šídák test; **Figure 3E**). (**Table S1** provides a detailed analysis of the spectro-temporal features of each syllable type across all ages tested in controls and *Magel2*^m+/p−^ pups).

In summary, we found that *Magel2*^m+/p−^ mice at P8 use simpler vocalizations that fall under the ‘flat’ and ‘short’ classifications instead of multicomponent USVs. These findings are in line with our previous results **(Figures 1-2)** demonstrating that the largest differences in vocal behavior occur in eight-day-old *Magel2*^m+/p−^ pups.

### Altered vocal repertoire of *Magel2*^m+/p−^ mice

As stated above, the vocal analysis pipeline outputs the probability for each USV to be classified as each of the eleven syllable types (*P*_1_, *P*_*2*_, *P*_*3*_, … *P*_11;_ **Figure 3A-B**). This distribution of probabilities allows the qualitative and quantitative comparison of the vocal classification among groups^21^. By considering the distribution of probabilities to classify each USV, it is possible to estimate how similar the vocal repertoire of one group of mice is to another group. To compare the vocal repertoire of mice across all ages studied, we used diffusion maps—a dimensionality reduction technique that decreases the number of dimensions of the probability distribution from eleven classes to three dimensions in a Euclidean space (**Figure 4A**)^21^. Using pairwise comparisons (**Figure 4B**), we estimated the similarity between the vocal repertoire of mice of different ages and genotypes. Using this method, we found that control pups at P6 and P8 (Cohen’s coefficient: κ = 0.99) and control pups at P10 and P12 (Cohen’s coefficient: κ = 0.95) display vocal repertoires that are similar to each other (**Figure 4C-D**). These two age groups (P6-P8 and P10-P12), however, present lower pairwise similarities when compared to each other with κ ranging from 0.67 to 0.77 (**Figure 4E**). These results suggest that the vocal repertoire of control pups undergoes significant changes between P8 and P10.

**Figure 4.**
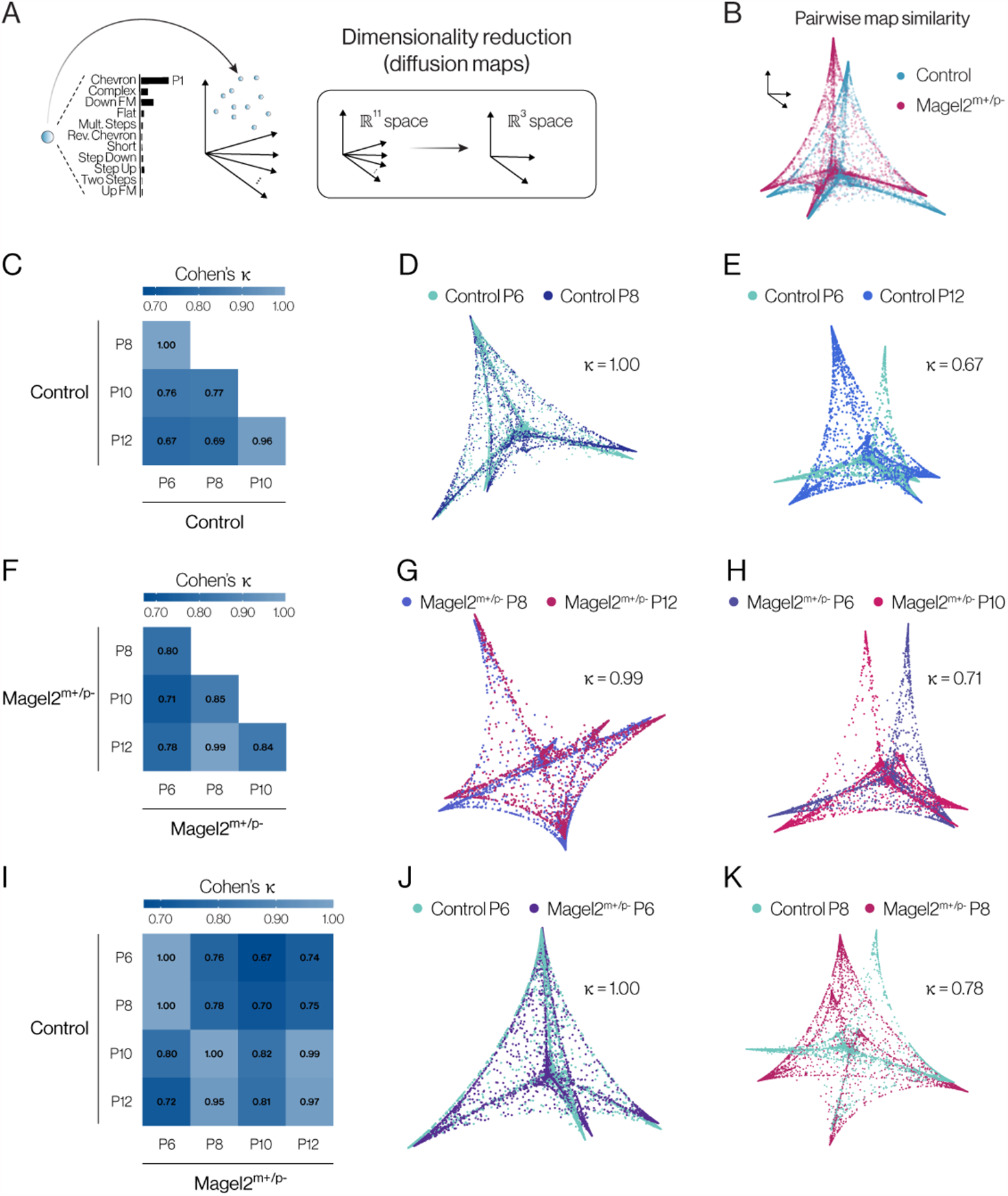
Analysis of the vocal repertoire of pups across ages. (**A**) Illustration of the output of the convolutional neural network, with a distribution of eleven probabilities for vocal classification (one probability for each of the eleven syllable types, with the highest probability defining the syllable type). Using diffusion maps, a dimensionality reduction technique, these eleven dimensions are reduced to three dimensions in the Euclidian space. (**B**) Illustration of a pairwise comparison of the vocal repertoire of pups using diffusion maps and 3D alignment of the manifolds (see methods for more details). (**D**) Comparison of the pairwise distance matrix between control pups at different ages using Cohen’s Kappa coefficient. (**D-E**) Examples of two pairwise comparisons with high and low alignment. (**F-H**) Similar to (**D-E**), but for *Magel2* deficient pups. (**I-K**) Similar to (C-H), but comparing control and *Magel2* deficient pups across ages. The sample sizes for control and *Magel2* deficient pups are: P6, n = 16 and 20; P8, n = 23 and 28; P10, n = 23 and 20; and P12, n = 30 and 28, respectively.

Next, we analyzed the same transitions in the vocal repertoire of *Magel2*^m+/p−^ pups. The comparison between the vocal repertoire of *Magel2*^m+/p−^ pups at P6 and P8 show lower pairwise similarity (κ = 0.80) compared to control pups (**Figure 4F**). In contrast to littermate controls, *Magel2*^m+/p−^ pups at P8 show a higher pairwise similarity with P10 (κ = 0.84) and P12 (κ = 0.99) pups (**Figure 4F-H)**. Finally, we directly compared the vocal repertoire of *Magel2*^m+/p−^ and control pups. *Magel2*^m+/p−^ pups, at P6, show high pairwise similarity when compared to controls at P6 (κ = 1.00) and P8 (κ = 1.00), but not at P10 (κ = 0.80) and P12 (κ = 0.72). At P8, *Magel2*^m+/p−^ pups show relatively low pairwise similarities with control pups at P6 (κ = 0.76) and P8 (κ = 0.78) but show high similarities with control pups at P10 (κ = 1.00) and P12 (κ = 0.95).

At P10, *Magel2*^m+/p−^ pups show relatively lower pairwise similarities with control pups at P6 (κ = 0.67) and P8 (κ = 0.70) than at P10 (κ = 0.82) and P12 (κ = 0.81). This pattern is more evident in P12 Magel*2*^m+/p−^ pups, which show lower pairwise similarities with control pups at P6 (κ = 0.74) and P8 (κ = 0.75) than at P10 (κ = 0.99) and P12 (κ = 0.97).

Altogether, these analyses suggest a different dynamic for the ontogeny of the vocal repertoire of *Magel2*^m+/p−^ compared to control pups—with *Magel2*^m+/p−^ pups at a younger age (i.e., P8) resembling control pups at an older age (i.e., P10-P12). Thus, the period of development between P8 and P10 seems to mark an important period for the effect of the maternally imprinted gene, *Magel2*, on the vocal behavior of the offspring.

### Dams prioritize their control offspring compared to *Magel2*^m+/p−^ in a retrieval test

The emission of vocalization by infants draws caregiver’s attention and care^12,13,15,17,22,23^ while genetically mute mouse pups are neglected by their dams^12^. Together, these observations led us to speculate that the deficiency of *Magel2* in pups, which alters vocal behavior, could lead to altered maternal behavior. In order to test this idea, we used a behavior assay to quantify maternal behavior towards their own control and *Magel2*^m+/p−^ offspring. In this assay, dams are first placed in the middle of a three-chamber apparatus that contains their home nest in the middle chamber (**Figure 5A**). After a period of acclimation, in the next stage, one pup of each genotype is placed at the opposite ends of the apparatus. The time (latency) to retrieve each pup back to the nest is then recorded (**Figure 5A**).

**Figure 5.**
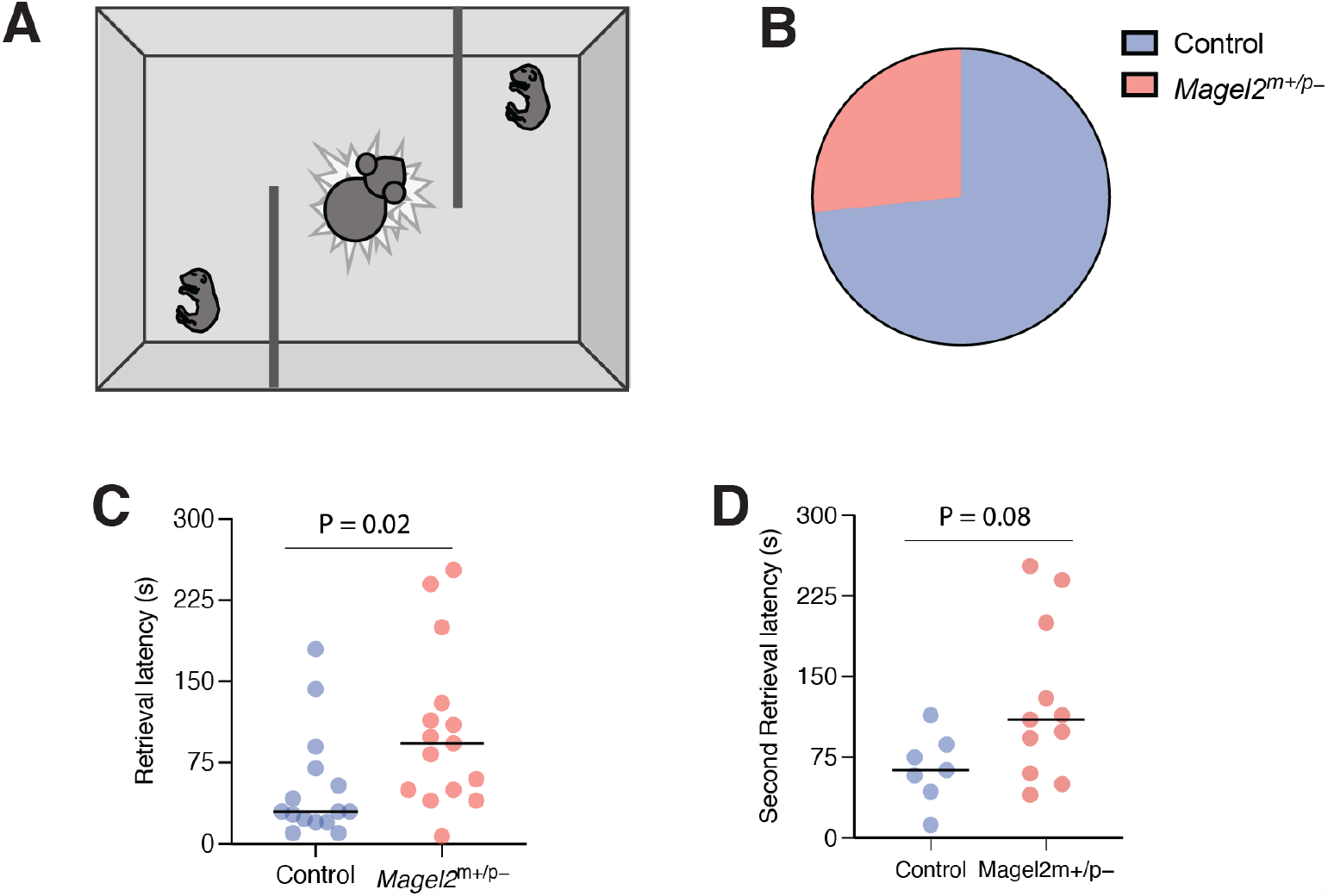
Maternal retrieval behavior is biased towards wildtype versus *Magel2* deficient pups. (**A**) Diagram of maternal retrieval protocol with P8 pups on opposite ends of the apparatus and mother at nest in the middle compartment. (**B**) Pie chart showing percentages of first retrieval of control and *Magel2* deficient pups from all trials. (**C**) Welch’s t-test comparing the retrieval latency of control and *Magel2* deficient pups. (**D**) Welch’s t-test comparing the latency for the retrieval of the second pup comparing *Magel2* deficient pups and wildtype controls that were picked second in the wildtype versus wildtype trial. Significant *P* values are shown in the graphs. The number *Magel2* deficient pups and controls trials are: P8, n = 15 and n = 7.

We initially evaluated whether dams show a preference to retrieve pups from one of the genotypes. This analysis shows that control pups were more likely to be retrieved first (11/15 trials; 73.3%) compared to *Magel2*^m+/p−^ pups (4/15 trials; 26.7%; **Figure 5B and Table S2**). Accordingly, on average, the latency to retrieve control pups was less than half of the latency to retrieve *Magel2*^m+/p−^ pups (control: 51.9 ± 12.9 s; *Magel2*^m+/p−^: 104.6 ± 19.1 s; *W* = 82, *P*_2-tail_ = 0.02, Wilcoxon test; **Figure 5C**). Because eleven out of fifteen *Magel2*^m+/p−^ pups were retrieved second, we also ran experiments placing wildtype control pups in both ends of the apparatus to compare the latency to retrieve the second pup in each of the experiments. Using this comparison, dams took twice as long to retrieve *Magel2*^m+/p−^ pups compared to control pups (control: 64.57 ± 12.3 s; *Magel2*^m+/p−^: 126.3 ± 22.23 s; *W* = 22, *P*_2-tail_ = 0.08, Mann-Whitney test; **Figure 5D**). In summary, the deficiency of Magel2 in offspring alters not only offspring behavior but also the behavior of the mother towards their own offspring.

## Discussion

In this study, we recorded and analyzed vocalizations from *Magel2* deficient pups and their wildtype littermates at postnatal days 6, 8, 10 and 12. Using custom-built software to automatically quantify vocalizations^21^, we counted the number of vocalizations and measured the spectro-temporal features of each vocalization, including its intensity, duration, bandwidth, mean frequency, maximum frequency, minimum frequency, and use of harmonic components. We also assigned a syllable type for each vocalization based on its morphological features in the time-frequency plane. We used further quantitative methods to analyze the vocal repertoire of mice across groups and ages. These methods shed light on discrete changes in the development of separation-induced vocalizations in *Magel2* deficient mice.

With regard to the vocal behavior of wildtype mice, our results demonstrate that the emission of separation-induced USVs gradually decreases from P6-P8 to P10-P12. This result agrees with previous studies, which show an inverse-U shape profile for separation-induced vocalizations in mice during the first two weeks of life ^15,29,31,32,38^. Based on the large number of vocalizations we analyzed, we also found that wildtype mice use simpler vocalizations at older ages (P10-P12) compared to younger ages (P6-P8). These findings in wildtype mice provide the basis for comparisons with *Magel2* deficient pups.

*Magel2* deficient pups show different dynamics for separation-induced vocalizations. At P6, these pups vocalize comparably to wildtype littermates, but at P8, their vocal number and features resemble wildtype littermates that are older (P10-P12). An explanation for these results is that *Magel2* deficient pups are less responsive to certain social cues. In socially isolated pups, therefore, the deprivation of these cues would not induce the behavior to the same degree as in wildtype pups. An alternative explanation for these results is that for *Magel2* deficient pups, vocal behavior does not have the same fitness value compared to wildtype pups. In the latter case, the behavior begins to change at younger ages due to the lack of reinforcement (maternal care). While it is difficult to prove these interpretations experimentally, the delayed latency for dams to retrieve *Magel2* deficient pups compared to wildtype littermates supports the idea that the deficiency of this imprinted gene impairs the fitness of the offspring. Moreover, the fact that *Magel2* deficient pups have lower body weight during early development further suggests a decrease in the fitness of these animals. Whether the change in vocal behavior and the decrease in body weight and maternal behavior are causally related warrants further investigation.

Interestingly, the deficiency of *Magel2* is not the only example of an imprinted gene affecting the emission of USVs by mouse pups. Consider, for example, previous studies of other imprinted genes. Deletion of the paternally inherited imprinted gene *Peg3* lowers vocal rate in mouse pups^39^. Conversely, deletion of the maternally inherited genes *Gabrb3* and *Ube3a* increases vocal rate^9^. Moreover, duplication of the paternal imprint loci on chromosome 15 increases the separation-induced vocalization of mouse pups^24^. Based on these findings, it is tempting to speculate that imprinted genes inherited from the father increase vocal rate while imprinted genes inherited from the mother decrease vocal rate. To test this generalization more formally, future studies will need to test the effect of *all* imprinted genes on the vocal behavior of the offspring.

P8 *Magel2*^m+/p−^ vocalize at a low rate, mainly emitting short and flat vocals similar to those of older pups. Moreover, when we estimate the similarity between the vocal repertoire of mutant and wildtype pups across ages tested, we find that P8 *Magel2*^m+/p−^ USVs are most similar to those produced by wildtype P10 and P12 pups. Interestingly, reduction in USV rate is associated to ASD, which is highly comorbid with MAGEL2^m+/p−^ patients^25^. Further studies are necessary to describe whether deviation in USV emission for *Magel2* deficient mice represents some kind of curve shift in the behavior. For example, it could be that the critical phase in which pups use separation-induced vocalization to gain maternal attention is shortened in offspring deficient for *Magel2*. Thus, our findings could suggest that alteration in the emission of USVs is part of a social behavior deficit in *Magel2*^m+/p−^ pups that continues into adulthood^26^.

In broader terms, we posit that our findings support the theory that genomic imprinting evolved to balance the cost of the phenotype for the offspring and for the mother, as well as to balance the best interests of mothers and fathers in altering offspring’s phenotype^3,29^. In the case of paternally inherited genes, the expression of these genes increases vocal behavior, thus, increasing maternal care and favoring the use of maternal resources, which is in the best interests of the father. Conversely, the loss of these paternally inherited genes decreases vocal behavior and, consequently, the demand for maternal care, thus, conserving maternal resources, which is in the best interests of the mother^2,5-7,27^. This theoretical framework is supported by our findings in a test that measures the latency to retrieve isolated pups back to the home nest, which demonstrate impaired maternal care towards Magel2 deficient pups compared to wildtype littermates^12,13,15,17,28^. As alluded above, future studies should test this theoretical view more directly by systematically investigating the effect of imprinted genes on the behavior of offspring and on the behavior of mothers towards their offspring. These efforts will help elucidate behavior phenotypes that occur in neurodevelopmental disorders as well as expand our understanding of the evolutionary adaptation of genomic imprinting.

## Acknowledgements

We thank the Dietrich Lab members for critical feedback on the project and the manuscript. Special thanks to: Delva Leão for her contribution to the data analysis; Jeremy Bober for helping with animal maintenance and care; as well as Antonio Fonseca who developed VocalMat software used for the analysis of vocal recordings. We thank David Gillich, Susan Andranovich, Valeria Krizsan, and Vickie Clark for administrative support. M.O.D. was supported by the National Institute of Mental Health of the National Institutes of Health under Award Number (R01MH125008) and by a grant of the Foundation for Prader-Willi Research. G.M.B.O. was supported by the HHMI Gilliam Fellowship. G.M.S. was supported by Coordenação de Aperfeiçoamento de Pessoal de Nível Superior - Brasil (CAPES). We thank David Bruin for copyediting the manuscript. The authors declare no conflict of interest.

## Author contributions

M.O.D and G.M.B.O. conceived the hypothesis, designed the study and wrote the manuscript. G.M.B.O. performed experiments. G.M.B.O. and G.M.S. analyzed and plotted the data. All authors read and edited the manuscript.

## Material and Methods

### Experimental models and subject details

All preweaning mice used in the experiments were 6 to 12 days old from both sexes (see table below for total number of recording used under each age specifying sex and genotype). Litters were provided from 9 separate breeding pairs. Separate litters were used for isolation and maternal retrieval tests. Details of litters tested found in **Table S1 and S2**. Dams used were 2 to 6 months old. To generate experimental pups, we used the following cross: *Magel2*^m+/p−^ (Jax #009062) dams bred with C57BL/6J (Jax #000664) males. Offspring from this cross were either *Magel2*^m+/p−^ or wildtype (*Magel2*^m+/p+^). All mice were kept in temperature- and humidity-controlled rooms, in a 12/12 hr. light/dark cycle, with lights on from 7:00 AM–7:00 PM. Studies took place during the light cycle. Food (Teklad 2018S, Envigo) and water were provided ad libitum. All procedures were approved by IACUC (Yale University).

### Behavior test

Pups from the same litter were placed individually in a soundproof chamber containing fresh bedding material ^19^. An UltraSoundGate Condenser Microphone CM 16 (Avisoft Bioacoustics, Berlin, Germany) was placed 10 cm above the recording chamber and connected to the UltraSoundGate 416 USGH device to record ultrasonic vocalizations. The recording sessions lasted 20 minutes. Four to eight chambers were recorded simultaneously. After testing, mice were placed back in their home cage with the dam. Pups were tested at postnatal days 6, 8, 10, and 12. Because pups that were naïve for the test—only tested at one specific age—show similar results as pups tested at multiple ages, we pooled all mice together for our analysis. Moreover, literature in the field have reported that repeated isolation testing does not affect infant vocal behavior^29,30^. Details of individual behavior measurements are found in **Table S1**.

### Vocalization analysis

Ultrasonic vocalizations were automatically extracted from audio recordings using a custom-built tool ^14^. In brief, audio recordings were converted from the time-domain to the frequency-domain using a 1024-point Fast Fourier Transform (FFT) through a 512-width hamming window with 50% overlap. Spectrograms were computed from the FFT and processed as images. Each pixel in the spectrogram corresponded to the intensity of each time-frequency component. Next, we applied a series of image-processing techniques (e.g., contrast enhancement, binarization, median filter, and morphological operations) to obtain segmentation of candidate vocalizations. A single spectrogram was generated for each candidate vocalization detected. Candidate vocalizations were classified as noise or real vocalization using a local median noise filter. The remaining vocalization candidates are further labeled under one of eleven call type classifications ^20^ using a Convolutional Neural Network (CNN), or as noise. The CNN was trained using a curated vocalization dataset, containing over 20,000 noise samples and 40,000 vocalization samples. Finally, the tool produces one spectrogram centralized on each vocalization for visual inspection, and a table (*xlsx* format) containing spectro-temporal features for each USV, such as time, duration, bandwidth, frequency, and intensity (minimum, mean, and maximum) values.

### Maternal Retrieval Test

The maternal preference test was performed in a three-chamber apparatus (65 × 42 × 23 cm) and comprised of three stages: Stage 1 – acclimation: the dam was allowed to explore the apparatus without the presence of pups for five minutes. In the middle of the apparatus, home cage nesting under an infrared igloo. Stage 2 – exploration: two P8 mice were placed on each side of the apparatus and the dam was allowed to explore the pups for five minutes. Stage 3 - preference: dam was recorded while she retrieved pups back to her home cage nesting. Groups were randomly alternated between both sides to avoid preference for one side of the chamber. Latency to retrieve each pup was timed from videos taken of the retrieval tests. Summary of results found in **Table S2**.

### Quantification and statistical analysis

Prism 8.0 or above was used to analyze data and plot figures. Shapiro-Wilk normality test was used to assess normal distribution of the data. To analyze differences in the use of harmonics, we used the non-parametric Mann-Whitney Test with Bonferroni correction to find statistically different effects. Comparison of diffusion maps were analyzed through pairwise comparison using Cohen’s kappa coefficient. Maternal retrieval latency for retrieval of each pup from the start of test used Wilcoxon test. Analysis of second retrieval latency from first retrieval was calculated using Welch’s t test. The rest of the data was analyzed using two-way ANOVA or mixed-effects analysis. Sidak’s multiple comparisons test was used to find post hoc differences among groups and to calculate the 95% confidence intervals to report effect size. In the text, values are provided as mean ± SEM. *P* < 0.05 was considered statistically significant and, when necessary and as described above, was corrected using Bonferroni’s method. Statistical data are provided in text and in the figures.

---

**Figure S1.**
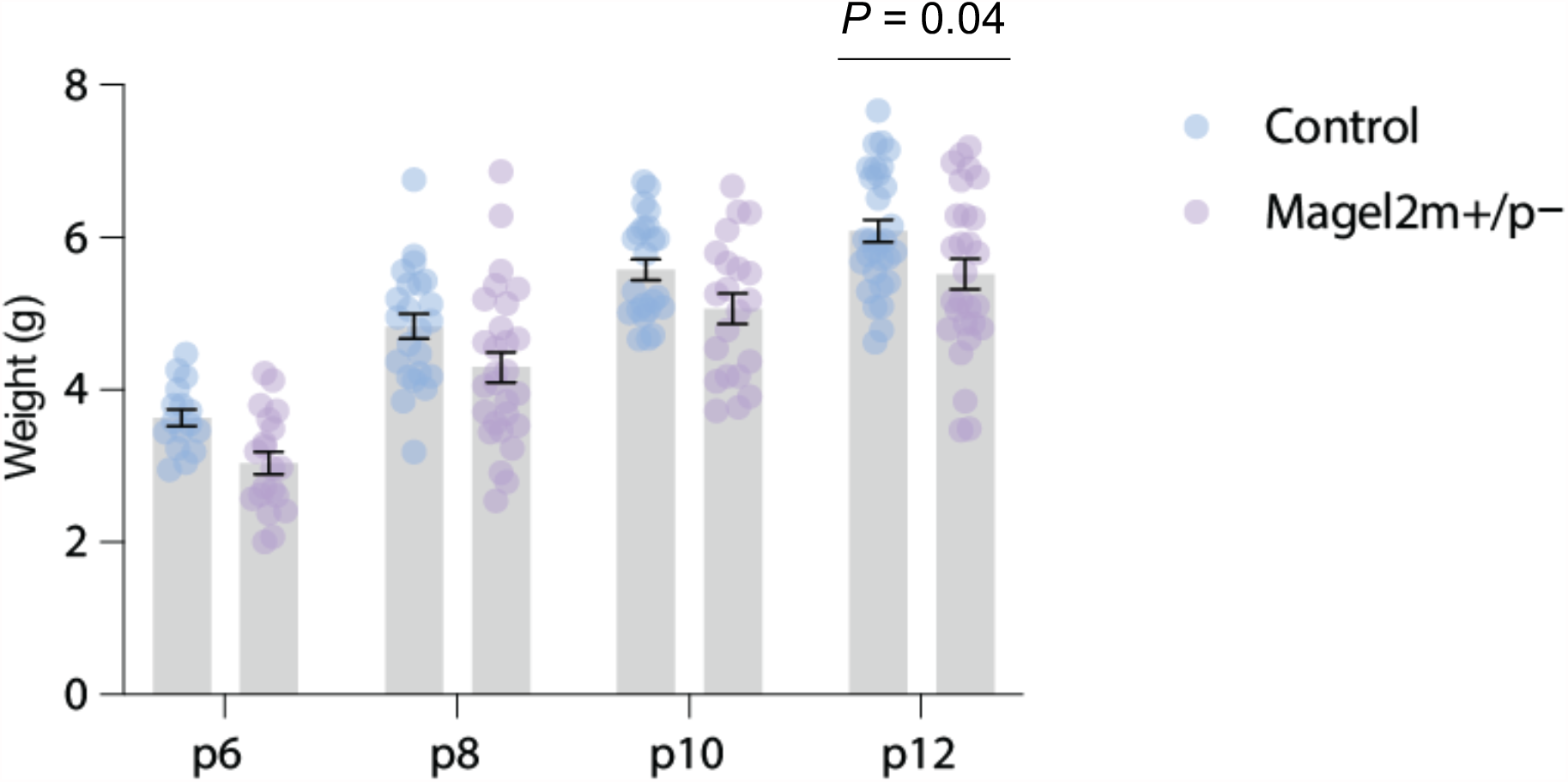
Body weight of pups across tested ages. Individual body weights of control (blue) and *Magel2* deficient (purple) littermates at P6, P8, P10, and P12. Line with error bars represent mean and SEM. Round symbols representing individual values. *P* values are provided in the main text. The sample sizes for control and *Magel2* deficient pups are: P6, n = 16 and 20; P8, n = 23 and 28; P10, n = 23 and 20; and P12, n = 30 and 28, respectively.

**Figure S2.**
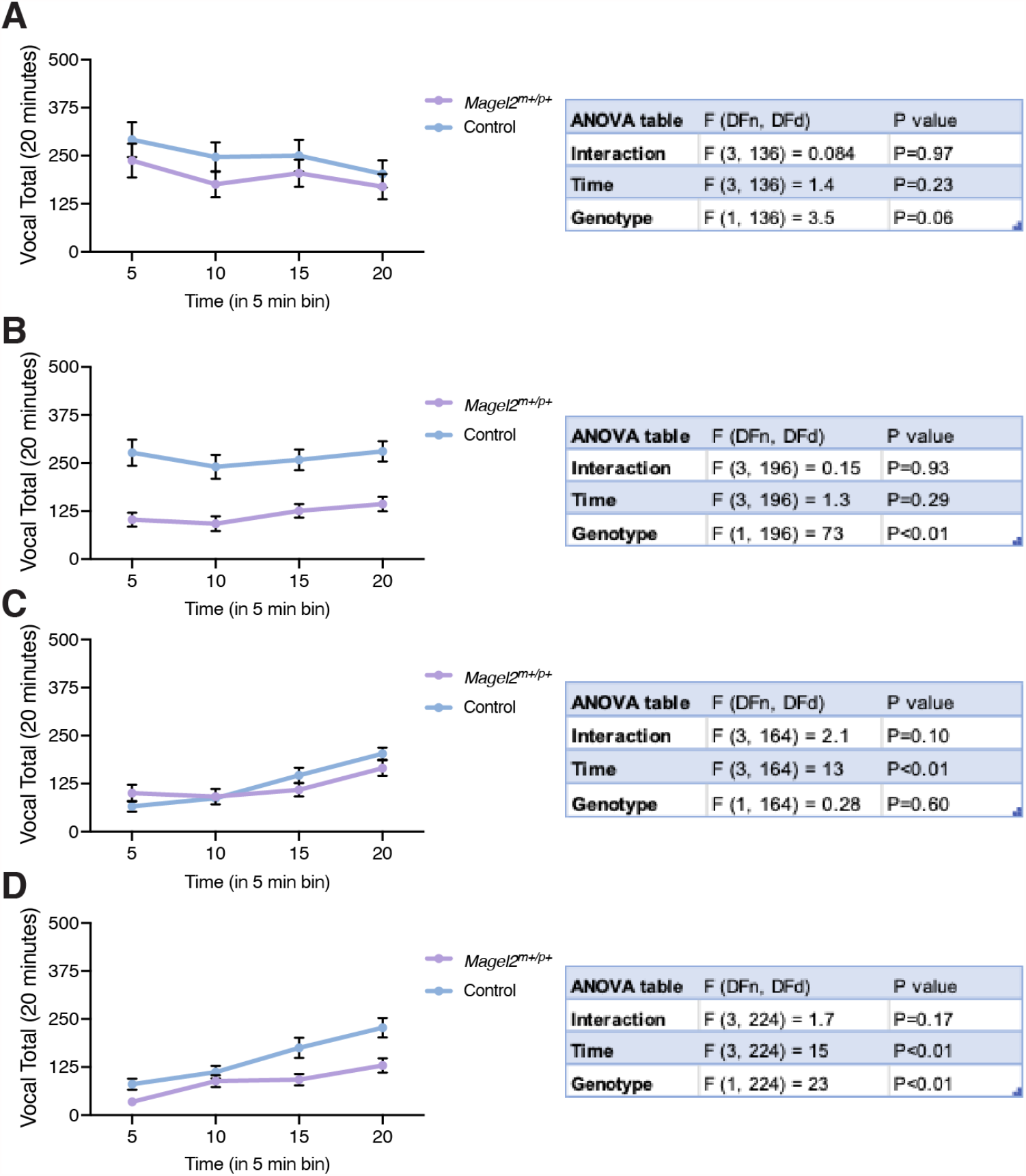
Vocal rate in 5-minute bins across ages. Average USV emission of control (blue) and *Magel2* deficient (purple) littermates at (A) P6, (B) P8, (C) P10, and (D) P12. Connecting lines trace USV means for each genotype across 5-minute bins from the 20-minute isolation recording. Line with error bars represent mean and SEM of weights. Round symbols represent mean values. 2way ANOVA results for each age are found on the tables on the right of each graph. The sample sizes for control and *Magel2* deficient pups are: P6, n = 16 and 20; P8, n = 23 and 28; P10, n = 23 and 20; and P12, n = 30 and 28, respectively.

**Figure S3.**
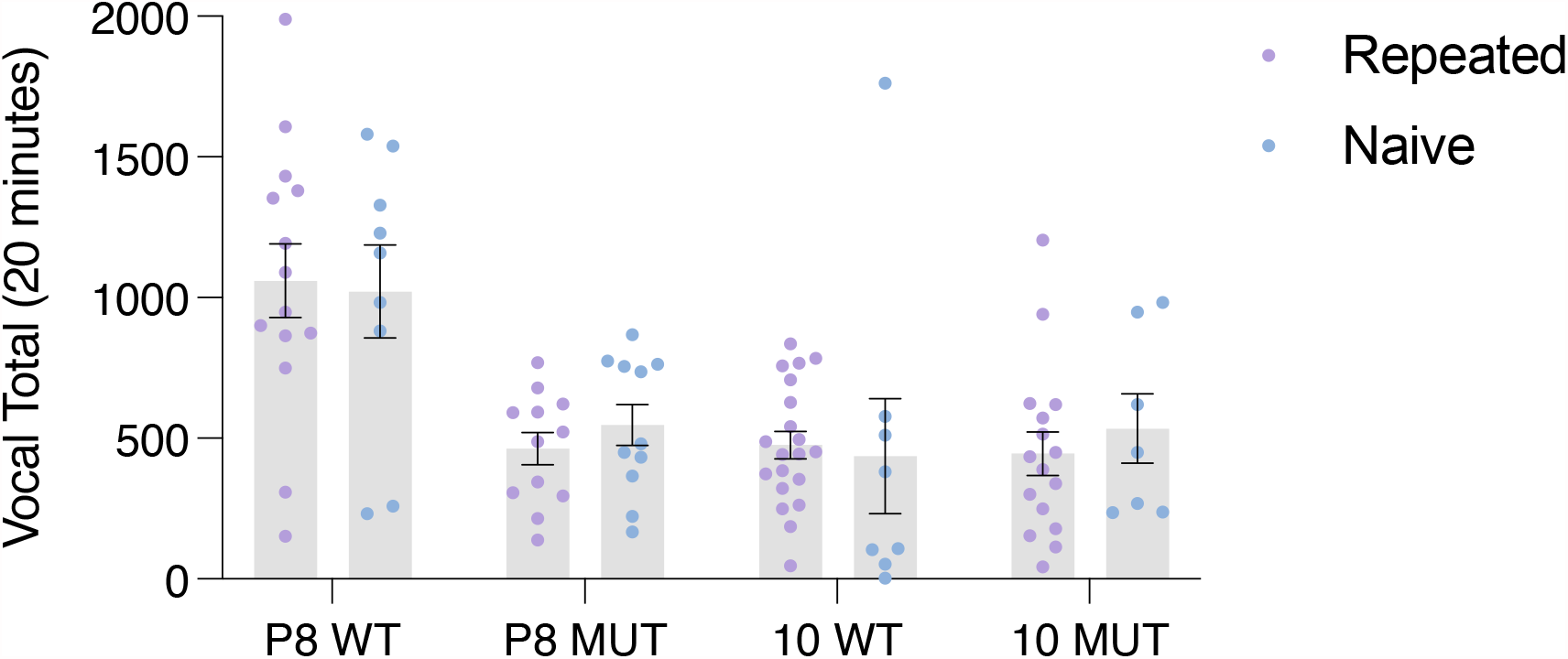
Rate of USVs in naïve versus experienced mouse pups. Average USV emission of pups tested repeatedly (purple) and naïve pups (blue). Bars represent USV means for each genotype and condition. Line with error bars represent mean and SEM. Round symbols represent rate for individual pups. Multiple Mann-Whitney test was used to compare rates. *P* values are the following: P8 WT = 0.36; P8 MUT = 0.46; P10 WT = 0.46; P10 MUT = 0.58.

